# The aging transcriptome and cellular landscape of the human lung in relation to SARS-CoV-2

**DOI:** 10.1101/2020.04.07.030684

**Authors:** Ryan D. Chow, Sidi Chen

## Abstract

Since the emergence of SARS-CoV-2 in December 2019, Coronavirus Disease-2019 (COVID-19) has rapidly spread across the globe. Epidemiologic studies have demonstrated that age is one of the strongest risk factors influencing the morbidity and mortality of COVID-19. Here, we interrogate the transcriptional features and cellular landscapes of the aging human lung through integrative analysis of bulk and single-cell transcriptomics. By intersecting these age-associated changes with experimental data on host interactions between SARS-CoV-2 or its relative SARS-CoV, we identify several age-associated factors that may contribute to the heightened severity of COVID-19 in older populations. We observed that age-associated gene expression and cell populations are significantly linked to the heightened severity of COVID-19 in older populations. The aging lung is characterized by increased vascular smooth muscle contraction, reduced mitochondrial activity, and decreased lipid metabolism. Lung epithelial cells, macrophages, and Th1 cells decrease in abundance with age, whereas fibroblasts, pericytes and CD4+ Tcm cells increase in abundance with age. Several age-associated genes have functional effects on SARS-CoV replication, and directly interact with the SARS-CoV-2 proteome. Interestingly, age-associated genes are heavily enriched among those induced or suppressed by SARS-CoV-2 infection. These analyses illuminate potential avenues for further studies on the relationship between the aging lung and COVID-19 pathogenesis, which may inform strategies to more effectively treat this disease.

## Main Text

Age is one of the strongest risk factors for severe outcomes among patients with COVID-19 ^1,2^. For instance, the case-fatality rate of patients 50-59 years old was reported to be 1.0% in Italy (as of March 17, 2020) and 1.3% in China (as of February 11, 2020) ^3,4^; in contrast, the case-fatality rate of patients ≥ 80 years old was 20.2% in Italy, and 14.8% in China during that same time frame (**Figure 1a**). Similarly, in the United States, the Center for Disease Control (CDC) estimated that from February 12 to March 16, 2020, the case-fatality rate of patients 55-64 years old was 1.4 – 2%, and 10.4 −27.3% for patients ≥ 85 years old ^5^ (**Figure 1a**). Furthermore, a recent analysis of Chinese CDC data revealed that while patients of all ages shared a similar probability of becoming infected by SARS-CoV-2, the causative agent of COVID-19 ^6–8^, the clinical manifestations of infection in children (< 18 years old) were less severe than in adults. With the exception of infants and younger children (< 1 years old and 1-5 years old, respectively), most children were asymptomatic or experienced mild illness^9^. Collectively, these observations indicate a strong association between age and COVID-19 morbidity and mortality. However, it must be emphasized that younger patients can still frequently contract the disease, possibly leading to hospitalization, ICU admission, or death (**Supplementary Figure 1**).

**Figure 1:**
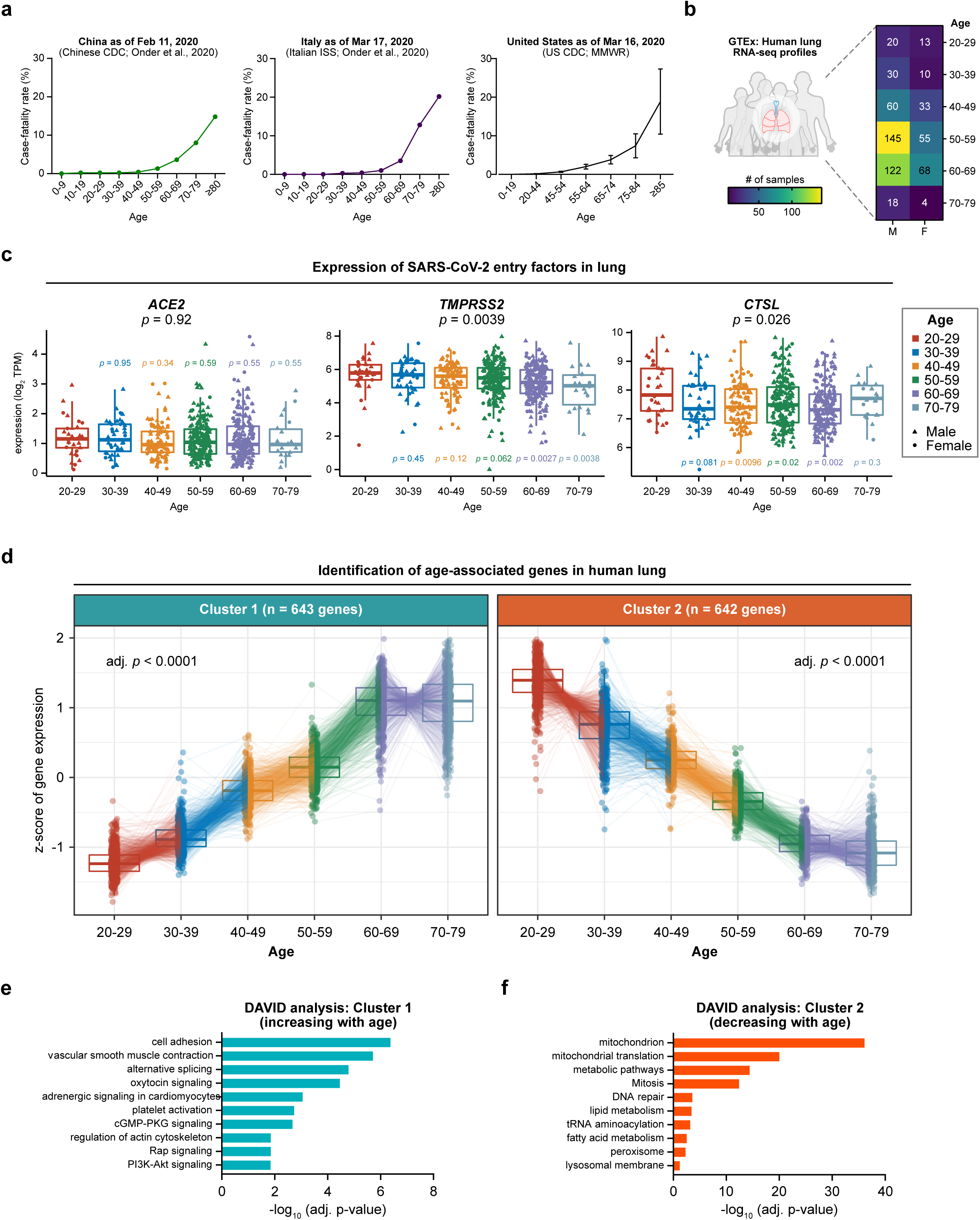
Identification of age-associated genes in the human lung. **a**. Case-fatality statistics in China, Italy, and the United States. Data are from the Chinese CDC (as of February 11, 2020), the Italian ISS (as of March 17, 2020), and the US CDC (as of March 16, 2020), respectively. Error bars for case-fatality in the United States represent lower and upper bounds of the estimated rates, given the preliminary nature of the data. **b**. Demographics of the human lung RNA-seq profiles in the GTEx dataset, detailed by sex and age group (total n = 578). **c**. Tukey boxplots detailing the expression of SARS-CoV-2 entry factors *ACE2, TMPRSS2*, and *CTSL* in the lung. Data are shown as log_2_ transcripts per million (TPM). Statistical significance of the expression variation across all age groups is shown on top of each plot (Kruskal-Wallis test). Pairwise comparisons of each age group to the 20-29 year-old group are also shown, color-coded by the age group (two-tailed Mann-Whitney test). **d**. Dot plot of age-associated genes in human lung. Cluster 1 contains genes that increase in expression with aging (left, n = 643), while cluster 2 contains genes that decrease in expression with aging (right, n = 642). Statistical significance was assessed by DESeq2 likelihood ratio test (adj. *p* < 0.0001). Data are shown in terms of median z-score, with lines connecting the same gene across different age groups. **e**. DAVID gene ontology and pathway analysis of Cluster 1 genes (increasing with age). **f**. DAVID gene ontology and pathway analysis of Cluster 2 genes (decreasing with age).

While the effects of age on COVID-19 are likely to be multifactorial, involving a complex blend of systemic and local factors, we hypothesized that tissue-intrinsic changes that occur with aging may offer valuable clues. To explore possible mechanisms by which age influences the clinical manifestations of SARS-CoV-2 infection, here we investigate the transcriptomic features and cellular landscape of the aging human lung in relation to SARS-CoV-2.

## Results

We focused our analysis on the Genotype-Tissue Expression (GTEx) project^10,11^, a comprehensive public resource of gene expression profiles from non-diseased tissue sites. As the lung is the primary organ affected by COVID-19, we specifically analyzed lung RNA-seq transcriptomes from donors of varying ages (20-79 years old) (**Figure 1b**). A total of 578 lung RNA-seq profiles were compiled, of which 31.66% were from women.

An initial hypothesis for why SARS-CoV-2 differentially affects patients of varying ages is that the expression of host factors essential for SARS-CoV-2 infection may increase with aging. To assess this possibility, we examined the gene expression of *ACE2*, which encodes the protein angiotensin-converting enzyme 2 that is coopted as the host receptor for SARS-CoV-2 ^8,12–15^. There was no statistically significant association between age and *ACE2* expression levels (Kruskal-Wallis test, *p* = 0.92) (**Figure 1c**). While ACE2 is the direct cell surface receptor for SARS-CoV-2, transmembrane serine protease 2 (TMPRSS2) and cathepsin L (CTSL) have been demonstrated to facilitate SARS-CoV-2 infection by priming the spike protein for host cell entry^14^. Expression of the corresponding genes *TMPRSS2* and *CTSL* was modestly associated with age (*p* = 0.0039 and *p* = 0.026, respectively) (**Figure 1c**), but *TMPRSS2* and *CTSL* expression levels tended to decrease with age. Of note, biological sex was not significantly associated with expression of *ACE2, TMPRSS2*, or *CTSL* (**Supplementary Figure 2a-c**), though it has been observed that males are more likely to be affected by COVID-19 than females^16–18^. Taken together, these data indicate that differential expression of SARS-CoV-2 host entry factors alone is unlikely to explain the relationship between age and severity of COVID-19 illness.

To discern the host cell types involved in COVID-19 entry, we turned to a single cell RNA-seq (scRNA-seq) dataset of 57,020 human lung cells from the Tissue Stability Cell Atlas^19^. In agreement with prior reports, analysis of the single cell lung transcriptomes revealed that alveolar type 2 (AT2) cells were comparatively enriched in *ACE2* and *TMPRSS2*-expressing cells ^20,21^ (**Supplementary Figure 3a-b**). However, *ACE2*-expressing cells represented only 1.69% of all AT2 cells, while 47.52% of AT2 cells expressed *TMPRSS2*. Alveolar type 1 (AT1) cells also showed detectable expression of *ACE2* and *TMPRSS2*, but at lower frequencies (0.39% and 26.70%). *CTSL* expression could be broadly detected in many different cell types including AT2 cells, but its expression was particularly pronounced in macrophages (**Supplementary Figure 3c**).

Since the expression of host entry factors *ACE2, TMPRSS2* and *CTSL* did not increase with age, we next sought to identify all age-associated genes expressed in the human lung (Methods). Using a likelihood-ratio test^22^, we pinpointed the genes for which age significantly impacts their expression. With a stringent cutoff of adjusted *p* < 0.0001, we identified two clusters of genes in which their expression progressively changes with age (**Figure 1d**). Cluster 1 is composed of 643 genes that increase in expression with age, while Cluster 2 contains 642 genes that decrease in expression with age. Gene ontology and pathway analysis of Cluster 1 genes (increasing with age) revealed significant enrichment for cell adhesion, vascular smooth muscle contraction, oxytocin signaling, and platelet activation, in addition to several other pathways (**Figure 1e**). These findings are consistent with known physiologic changes of aging, including decreased pulmonary compliance^23^, and heightened risk for thrombotic diseases^24^. Of note, deregulation of the renin-angiotensin system has been implicated in the pathogenesis of acute lung injury induced by SARS-CoV ^25,26^, the closely related coronavirus responsible for the SARS epidemic of 2002-2003 ^27^. In mice, SARS-CoV infection downregulates ACE2, leading to disinhibition of angiotensin II production by ACE ^25,26,28–30^ and subsequent vasoconstriction. A possible hypothesis is that increased baseline vascular smooth muscle contraction in older patients may predispose the development of acute respiratory distress syndrome (ARDS) in the setting of SARS-CoV and/or SARS-CoV-2 infection. In line with this hypothesis, recent analyses of COVID-19 cohorts in China and Italy have found that patients with hypertension were more likely to develop ARDS ^17^, require ICU admission ^31^, and die from the disease ^32^, though we note that correlative epidemiologic studies do not necessarily demonstrate causality.

Cluster 2 genes (decreasing with age) were significantly enriched for mitochondrion, mitochondrial translation, metabolic pathways, and mitosis, among other pathways (**Figure 1f**), which is consistent with prior observations of progressive mitochondrial dysfunction with aging^33–35^. Of note, Cluster 2 was also enriched for genes involved in lipid metabolism, fatty acid metabolism, peroxisome, and lysosomal membranes. Age-associated alterations in lipid metabolism could impact SARS-CoV-2 infection, as SARS-CoV can enter cells through cholesterol-rich lipid rafts ^36–39^. Similarly, age-associated alterations in lysosomes could influence late endocytic viral entry, as the protease cathepsin L cleaves SARS-CoV spike proteins from within lysosomes ^40,41^.

Having compiled a high-confidence set of age-associated genes, we sought to identify the lung cell types that normally express these genes, using the human lung single cell transcriptomics dataset from the Tissue Stability Cell Atlas^19^. By examining the scaled percentage of expressing cells within each cell subset, we identified age-associated genes predominantly enriched in different cell types. Cell types with highly enriched expression for certain Cluster 1 genes (increasing with age) included fibroblasts, muscle cells, and lymph vessels (**Figure 2a**). In contrast, cell types with highly enriched expression for certain Cluster 2 genes (decreasing with age) included macrophages, dividing dendritic cells (DCs)/monocytes, and AT2 cells (**Figure 2b**). Similar results were found using an independent human lung scRNA-seq dataset from the Human Lung Cell Atlas (**Supplementary Figure 4a-b**)^42^. Examining the muscle-enriched genes that increased in expression with age, gene ontology analysis revealed enrichment for vascular smooth muscle contraction, cGMP-PKG signaling, Z-disc, and actin cytoskeleton, among other pathways **(Figure 2c**). As for the AT2-enriched genes that decreased in expression with age, gene ontology analysis revealed enrichment for metabolic pathways, biosynthesis of antibiotics, lipid metabolism, extracellular exosome, and mitochondrial matrix (**Figure 2d**). A subset of these enriched gene ontologies had also been identified by the bulk RNA-seq analysis (**Figure 1e-f**). Thus, integrative analysis of bulk and single-cell transcriptomes revealed that many of the age-associated transcriptional changes in human lung can be mapped to specific cell subpopulations, suggesting that the overall abundance of these cell types, their transcriptional status, or both, may be altered with aging.

**Figure 2:**
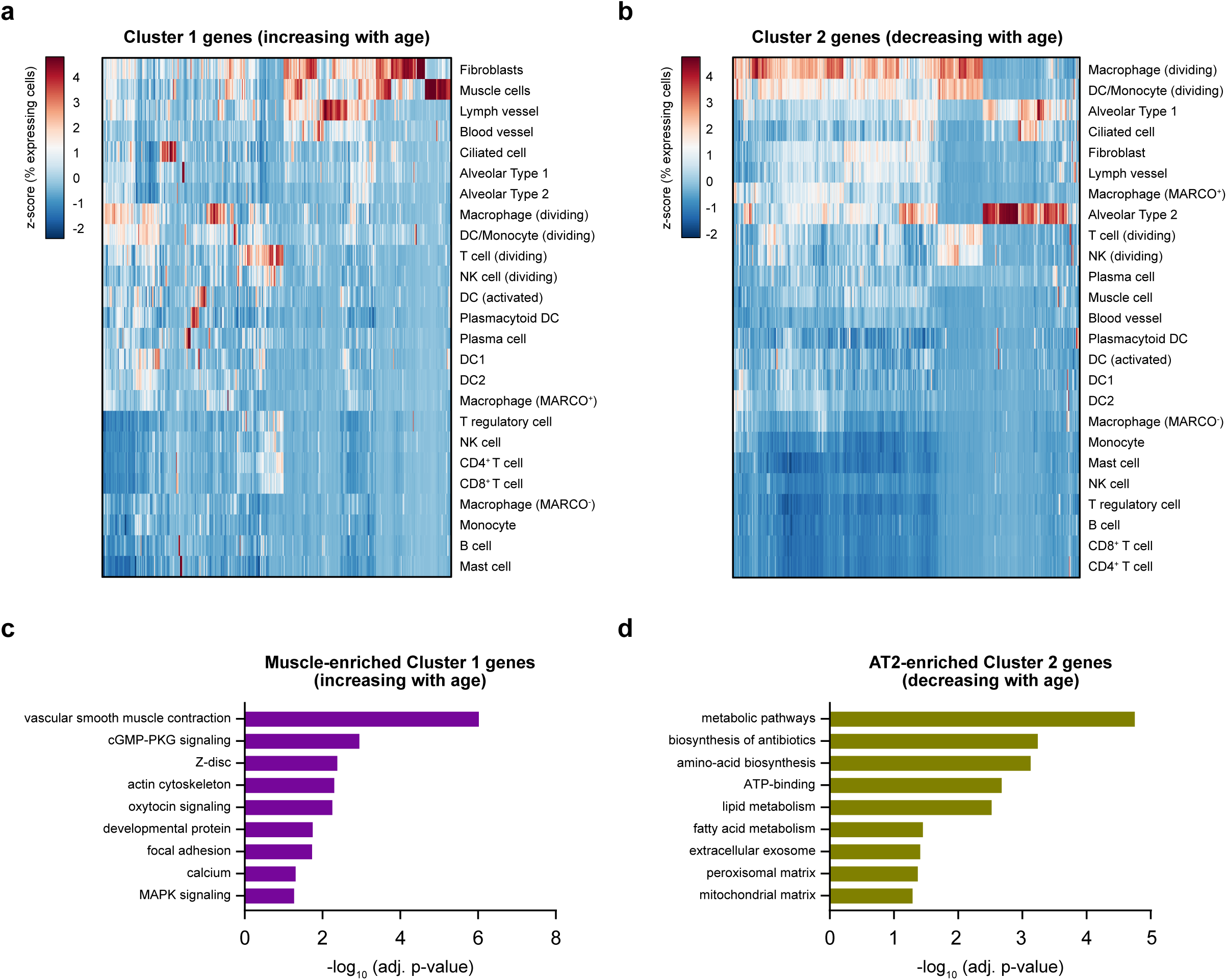
Lung single-cell transcriptomics pinpoints cell type-specific expression of age-associated genes. **a**. Heatmap showing the percentage of cells expressing each of the Cluster 1 genes (increasing with age), scaled by gene across the different cell types. Data are from the Tissue Stability Cell Atlas. **b**. Heatmap showing the percentage of cells expressing each of the Cluster 2 genes (decreasing with age), scaled by gene across the different cell types. Data are from the Tissue Stability Cell Atlas. **c**. DAVID gene ontology and pathway analysis of Cluster 1 age-associated genes that exhibit enriched expression in muscle cells. **d**. DAVID gene ontology and pathway analysis of Cluster 2 age-associated genes that exhibit enriched expression in alveolar type 2 (AT2) cells.

As the pathophysiology of viral-induced ARDS involves an intricate interplay of diverse cell types, most notably the immune system^43,44^, aging-associated shifts in the lung cellular milieu^23^ could contribute an important dimension to the relationship between age and risk of ARDS in patients with COVID-19 ^31^. To investigate the cellular landscape of the aging lung, we applied a gene signature-based approach^45^ to infer the enrichment of different cell types from the bulk RNA-seq profiles. Since bulk RNA-seq measures the average expression of genes within a cell population, such datasets will reflect the relative proportions of the cell types that comprised the input population, though with the caveat that cell types can have overlapping expression profiles and such profiles may be altered in response to stimuli. Using this approach, we identified age-associated alterations in the enrichment scores of several cell types (**Figure 3a**). Whereas epithelial cells decreased with age, fibroblasts increased with age (**Figure 3b**). This finding is consistent with the progressive loss of lung parenchyma due to reduced regenerative capacity of the aging lung^46^, as well as the increased risk for diseases such as chronic obstructive pulmonary disease and pulmonary fibrosis^47^. In addition, these results are concordant with the findings from analysis of human lung single-cell transcriptomes (**Figure 2a-b**).

**Figure 3:**
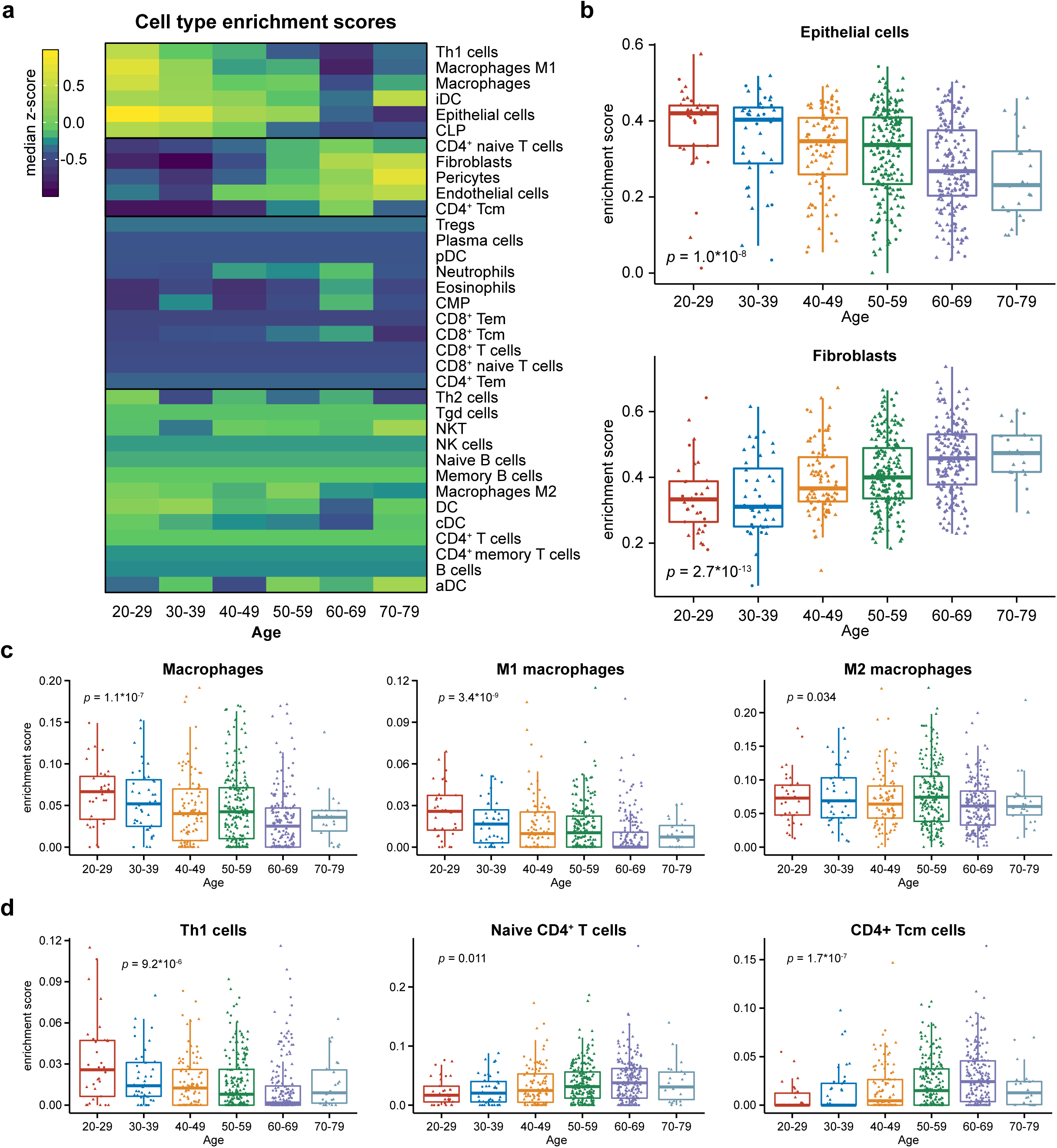
The evolving cellular landscape of the aging human lung. **a**. Heatmap of cell type enrichment scores in the human lung across different age groups. Data are expressed as median z-scores within each subpopulation. **b**. Tukey boxplots detailing the enrichment scores of epithelial cells (top) and fibroblasts (bottom) in the lung across age groups. Statistical significance of age-associated variation was assessed by Kruskal-Wallis test. **c**. Tukey boxplots detailing the enrichment scores of macrophages, M1 macrophages, and M2 macrophages in the lung across age groups. Statistical significance of age-associated variation was assessed by Kruskal-Wallis test. **d**. Tukey boxplots detailing the enrichment scores of Th1 cells, naive CD4^+^ T cells, and CD4^+^ Tcm cells in the lung across age groups. Statistical significance of age-associated variation was assessed by Kruskal-Wallis test.

Among the innate immune cell populations, the enrichment scores of total macrophages were inversely associated with age (**Figure 3c**). Macrophages are major drivers of innate immune responses in the lung, acting as first-responders against diverse respiratory infections^48^. Thus, the age-associated decrease in macrophage abundance may be a possible factor related to the greater severity of lung pathology in patients with COVID-19. Although macrophage accumulation is often associated with the pathologic inflammation of viral ARDS ^49,50^, pulmonary macrophages can act to limit the duration and severity of infection by efficiently phagocytosing dead infected cells and released virions ^51–53^. Notably, macrophages infected with SARS-CoV have been found to abort the replication cycle of the virus^54,55^, further supporting their role in antiviral responses. However, macrophages may suppress antiviral adaptive immune responses ^48,56^, inhibiting viral clearance in mouse models of SARS-CoV infection ^57^. In aggregate, these prior reports suggest that the precise role of lung macrophages in SARS-CoV-2 pathophysiology is likely context-dependent. It is also plausible that the increased numbers of macrophages are not the primary distinction between young and old patients, but rather the functional status of the macrophages. In line with this, we observed that the age-associated changes in macrophages were specifically attributed to the pro-inflammatory M1 macrophage subset but not the immunoregulatory M2 subset ^58^ (**Figure 3c**), though this binary classification scheme represents an oversimplification of macrophage function. Nevertheless, elucidating the consequences of age-associated changes in lung macrophages may reveal insights into the differential outcomes of older patients with COVID-19. Further studies are needed to investigate whether macrophages or other innate immune cells respond to SARS-CoV-2 infection, and how their numbers or function may change with aging.

Among the adaptive immune cell populations, we observed that Th1 cells and CD4^+^ Tcm cells trended in opposite directions with aging (**Figure 3d**). While the lungs of younger donors were enriched for Th1 cells, they were comparatively depleted for CD4^+^ Tcm cells; the inverse was true in the lungs of older donors. Of note, mouse models of SARS-CoV infection have indicated important roles for CD4^+^ T cell responses in viral clearance ^59,60^. Additionally, Th1 cells are responsive to SARS-CoV vaccines ^61^ and promote macrophage activation against viruses ^62^. It is therefore possible that age-associated shifts in CD4^+^ T cell subtypes within the lung may influence the subsequent host immune response in response to coronavirus infection. However, future studies will be needed to determine the role of Th1 cells and other adaptive immune cells in the response to SARS-CoV-2, and how these dynamics may change with aging.

We next explored the roles of lung age-associated genes in host responses to viral infection. Since functional screening data with SARS-CoV-2 has not yet been described (as of March 30, 2020), we instead searched for data on SARS-CoV. While these two viruses belong to the same genus (*Betacoronaviridae*) and are conserved to some extent ^8^, they are nevertheless two distinct viruses with different epidemiological features, indicating unique virology and host biology. Therefore, data from experiments performed with SARS-CoV must be interpreted with caution.

We reassessed the results from a prior *in vitro* siRNA screen of host factors involved in SARS-CoV infection ^63^. In this kinase-focused screen, 130 factors were determined to have a significant effect on SARS-CoV replication. Notably, 11 of the 130 factors exhibited age-associated gene expression patterns (**Figure 4a**), with 4 genes in Cluster 1 (increasing with age) and 7 genes in Cluster 2 (decreasing with age). The 4 genes in Cluster 1 were all associated with increased SARS infectivity upon siRNA knockdown; these genes included *CLK1, AKAP6, ALPK2*, and *ITK*. Paradoxically, while knockdown of *CLK1* was associated with increased SARS infectivity, cell viability was also found to be increased (**Figure 4b**). Of the 7 genes in Cluster 2, 6 were associated with increased SARS infectivity and reduced cell viability upon siRNA knockdown (*AURKB, CDKL2, PDIK1L, CDKN3, MST1R*, and *ADK*). Age-related downregulation of these 6 factors could be related to the increased severity of illness in older patients. However, we emphasize that until rigorous follow-up experiments are performed with SARS-CoV-2, the therapeutic potential of targeting these factors in patients with COVID-19 is unknown.

**Figure 4:**
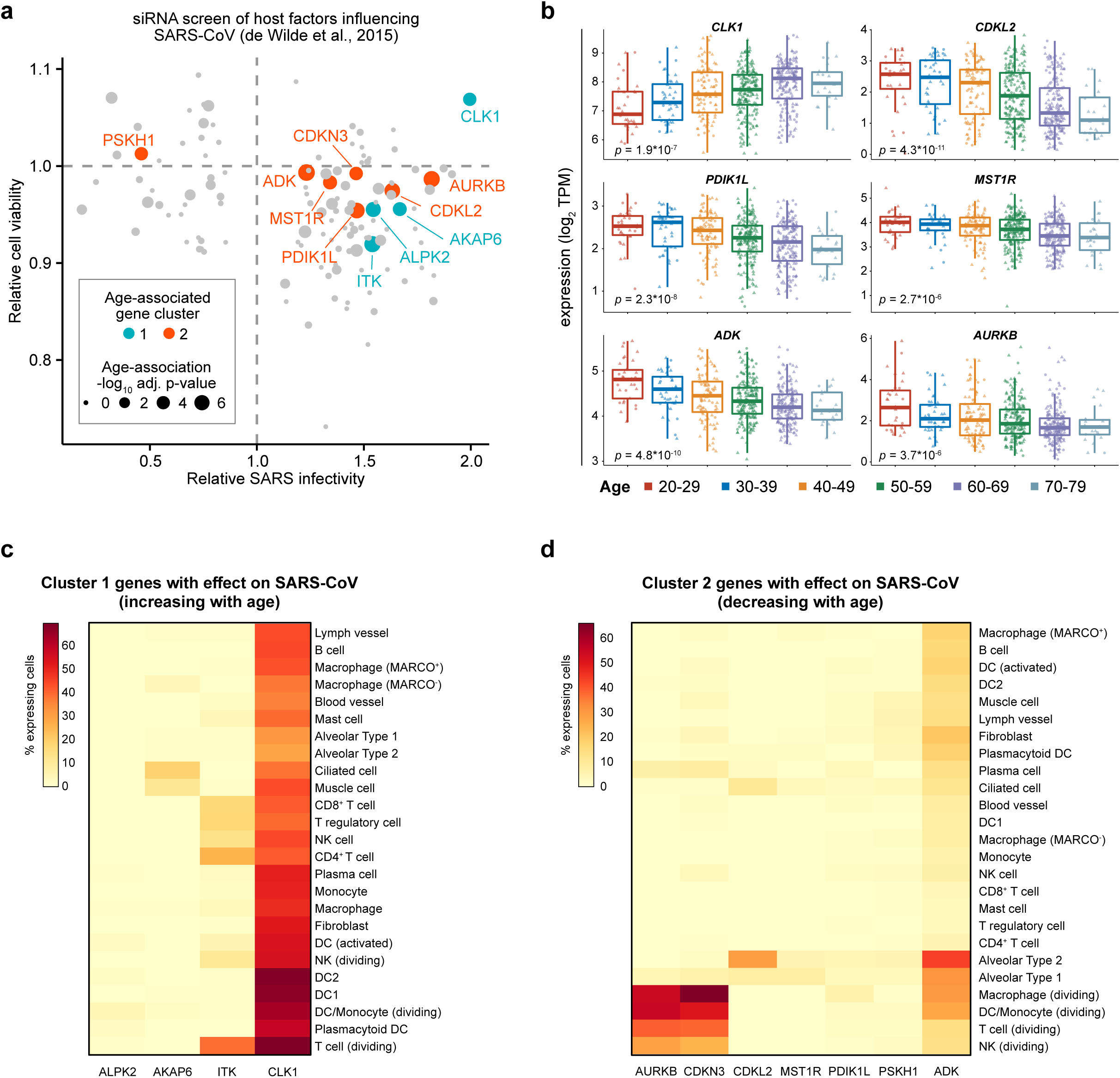
Age-associated genes in human lung influence SARS-CoV replication. **a**. Scatter plot of data from a published siRNA screen of host factors that influence SARS-CoV pathogenesis (from de Wilde et al., 2015). Data are shown in terms of relative SARS-CoV infectivity and relative cell viability upon perturbation of various host factors. Values >1.0 for relative SARS-CoV infectivity suggest that knockdown of the target gene promotes infection. Values >1.0 for relative cell viability suggest that knockdown of the target gene promotes cell survival. The size of each point is scaled by the statistical strength of the association between expression of the gene and aging. Genes identified in Cluster 1 (increasing with age) and Cluster 2 (decreasing with age) are additionally color-coded (blue and orange, respectively). **b**. Tukey boxplots detailing the expression of select age-associated genes with potential roles in SARS-CoV pathogenesis, highlighted in **(d)**. Data are shown as log_2_ transcripts per million (TPM) for *CLK1, CDKL2, PDIK1L, MST1R, ADK*, and *AURKB*. Statistical significance of the expression variation across all age groups was assessed by Kruskal-Wallis test. **c**. Heatmap showing the percentage of cells expressing each of the Cluster 1 genes (increasing with age) with an effect on SARS-CoV, as highlighted in **(a)**. Data are from the Tissue Stability Cell Atlas. **d**. Heatmap showing the percentage of cells expressing each of the Cluster 2 genes (decreasing with age) with an effect on SARS-CoV, as highlighted in **(a)**. Data are from the Tissue Stability Cell Atlas.

Using the human lung scRNA-seq data, we then determined which cell types predominantly express these host factors. Of the 4 genes in Cluster 1 that had a significant impact on SARS-CoV replication, *CLK1* was universally expressed, while ALPK2 expression was rarely detected (**Figure 4c, Supplementary Figure 5a**). *ITK* was preferentially expressed in lymphocytic populations, and *AKAP6* was most frequently expressed in ciliated cells and muscle cells. Of the 7 overlapping genes in Cluster 2, *AURKB* and *CDKN3* were predominantly expressed in proliferating immune cell populations, such as macrophages, DCs/monocytes, T cells, and NK cells (**Figure 4d, Supplementary Figure 5b**). *MST1R, PDIK1L*, and *PSKH1* were infrequently expressed, through their expression was detected in a portion of AT2 cells (5.54%, 5.30%, and 5.47%, respectively). Finally, *ADK* and *CDKL2* exhibited preferential enrichment in AT2 cells (51.40% and 34.43%). In aggregate, these analyses showed that the age-associated genes with functional roles in SARS-CoV are expressed in specific cell types of the human lung.

We then investigated whether age-associated genes in the human lung interact with proteins encoded by SARS-CoV-2. A recent study interrogated the human host factors that interact with 27 different SARS-CoV-2 proteins ^64^, revealing the SARS-CoV-2 : Human protein interactome in cell lines expressing recombinant SARS-CoV-2 proteins. By cross-referencing the interacting host factors with the set of age-associated genes, we identified 20 factors at the intersection (**Figure 5a**). 4 of these genes showed an increase in expression with age (i.e. Cluster 1 genes), while 16 decreased in expression with age (Cluster 2 genes). Mapping these factors to their interacting SARS-CoV-2 proteins, we noted that the age-associated host factors which interact with M, Nsp13, Nsp1, Nsp7, Nsp8, Orf3a, Orf8, Orf9c, and Orf10 proteins generally decrease in expression with aging (**Figure 5b**). However, a notable exception was Nsp12, as the age-associated host-factors that interact with Nsp12 both showed increased expression with aging (*CRTC3* and *MYCBP2*) (**Figure 5c**). Nsp12 encodes for the primary RNA-dependent RNA polymerase (RdRp) of SARS-CoV-2, and is a prime target for developing therapies against COVID-19. The observation that *CRTC3* and *MYCBP2* increase in expression with aging is intriguing, as these genes may be related to the activity of Nsp12/RdRp in host cells. Of note, MYCBP2 is a known repressor of cAMP signaling ^65,66^, and cAMP signaling potently inhibits contraction of airway smooth muscle cells ^67^. Thus, age-associated increases in *MYCBP2* could promote smooth muscle contraction, which is concordant with our analyses on age-associated gene signatures in the lung (**Figure 1e**). MYCBP2 might possibly contribute to COVID-19 pathology by not only interacting with SARS-CoV-2 RdRp, but also through its normal physiologic role in promoting smooth muscle contraction.

**Figure 5:**
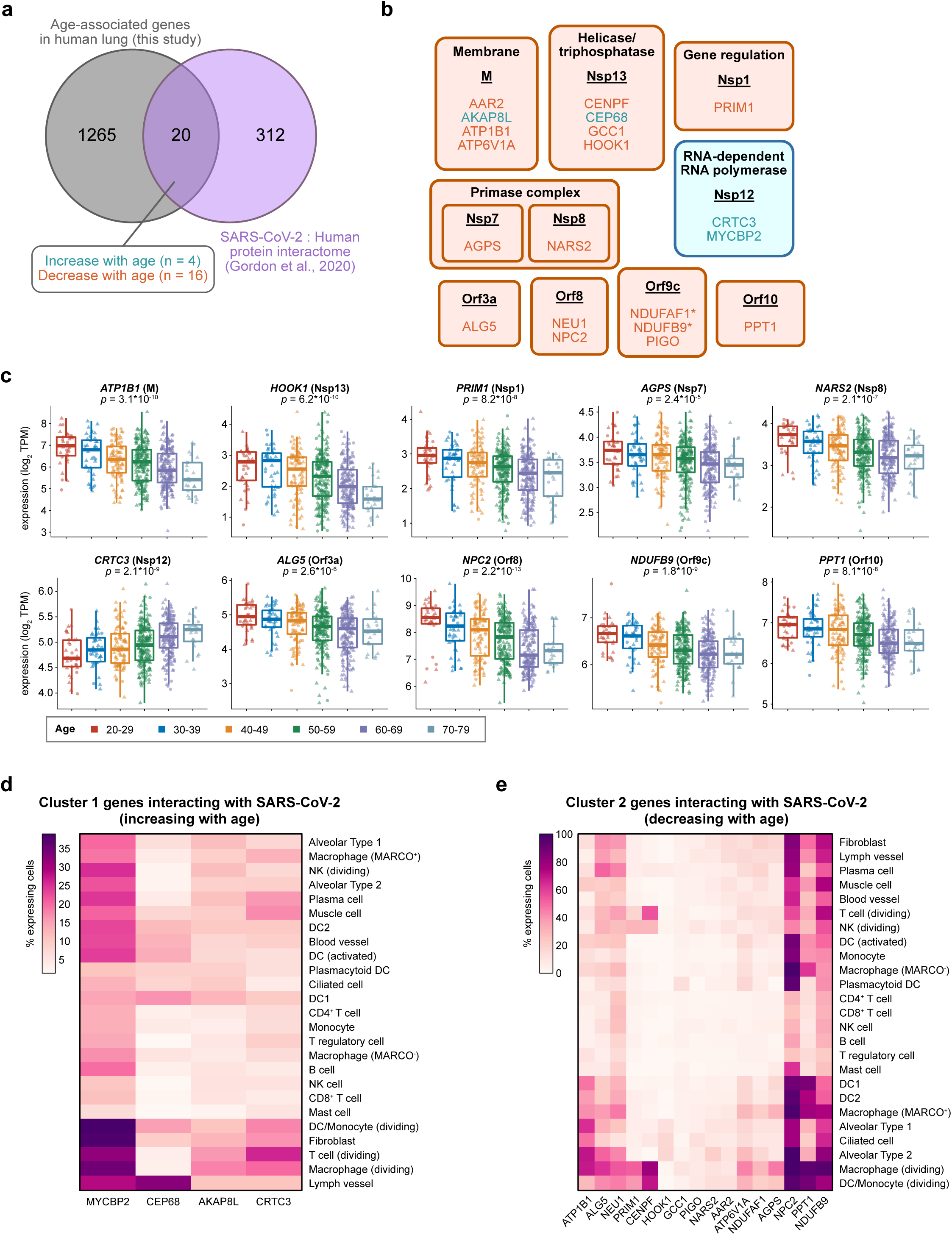
Age-associated genes in human lung interact with SARS-CoV-2 proteins. **a**. Venn diagram of the intersection between age-associated genes in human lung and the SARS-CoV-2 : Human protein interactome (Gordon et al., 2020). Of the 20 age-associated genes that were found to also interact with SARS-CoV-2, 4 of them increased in expression with age, while 16 decreased with age. **b**. Age-associated genes in human lung and their interaction with SARS-CoV-2 proteins, where each block contains a SARS-CoV-2 protein (underlined) and its interacting age-associated factors. Blocks are colored by the dominant directionality of the age association (orange, decreasing with age; blue, increasing with age). Gene targets with already approved drugs, investigational new drugs, or preclinical molecules are additionally denoted with an asterisk. **c**. Tukey boxplots detailing the expression of select age-associated genes that interact with SARS-CoV-2 proteins, highlighted in **(b)**. Data are shown as log_2_ transcripts per million (TPM) for *ATP1B1, HOOK1, PRIM1, AGPS, NARS2, CRTC3, ALG5, NPC2, NDUFB9*, and *PPT1*. The SARS-CoV-2 interacting protein is also annotated in parentheses. Statistical significance of the expression variation across all age groups was assessed by Kruskal-Wallis test. **d**. Heatmap showing the percentage of cells expressing each of the Cluster 1 genes (increasing with age) that interact with SARS-CoV-2 proteins, as highlighted in **(b)**. Data are from the Tissue Stability Cell Atlas. **e**. Heatmap showing the percentage of cells expressing each of the Cluster 2 genes (decreasing with age) that interact with SARS-CoV-2 proteins, as highlighted in **(b)**. Data are from the Tissue Stability Cell Atlas.

To assess the cell type-specific expression patterns of these various factors, we further analyzed the lung scRNA-seq data. Of the SARS-CoV-2 interacting genes that increase in expression with age, *MYCBP2* was frequently expressed across several populations, particularly proliferating immune populations (DC/monocyte, T cells, macrophages), muscle cells, fibroblasts, and lymph vessels (**Figure 5d**). *MYCBP2* was also expressed in 21.86% of AT2 cells. *CEP68* was preferentially expressed in lymph vessels, while *AKAP8L* and *CRTC3* showed relatively uniform expression frequencies across cell types, including a fraction of AT2 cells (8.53% and 9.07% expressing cells, respectively). Of the SARS-CoV-2 interacting genes that decrease in expression age, *NPC2* and *NDUFB9* were broadly expressed in many cell types, including AT2 cells (99.96% and 81.91%, respectively) (**Figure 5e**). AT2 cells also frequently expressed *ATP1B1, ALG5, NEU1*, and *ATP6V1A* (70.21%, 50.64%, 43.24%, and 27.78%). Analysis of the independent Human Lung Cell Atlas dataset revealed similar conclusions (**Supplementary Figure 6a-b**). Together, these analyses highlight specific age-associated factors that interact with the SARS-CoV-2 proteome, in the context of the lung cell types in which these factors are normally expressed.

Finally, we assessed whether SARS-CoV-2 infection directly alters the expression of lung age-associated genes. A recent study profiled the *in vitro* transcriptional changes associated with SARS-CoV-2 infection in different human lung cell lines ^68^. We specifically focused on the data from A549 lung cancer cells, A549 cells transduced with an ACE2 expression vector (A549-ACE2), and Calu-3 lung cancer cells. Several age-associated genes were found to be differentially expressed upon SARS-CoV-2 infection (**Figure 6a-c**). Of note, the overlap between lung age-associated genes and SARS-CoV-2 regulated genes was statistically significant across all 3 cell lines (**Figure 6d-f**), suggesting a degree of similarity between the transcriptional changes associated with aging and with SARS-CoV-2 infection. Among the age-associated genes that were induced by SARS-CoV-2 infection, the majority of these genes increase in expression with age (Cluster 1) (**Figure 6g-i**). Conversely, among the age-associated genes that were repressed by SARS-CoV-2 infection, most of these genes decrease in expression with age (Cluster 2). Of note, the directionality of SARS-CoV-2 regulation (induced or repressed) and the directionality of age-association (increase or decrease with age) were significantly associated across all 3 cell lines (**Figure 6g-i**).

**Figure 6:**
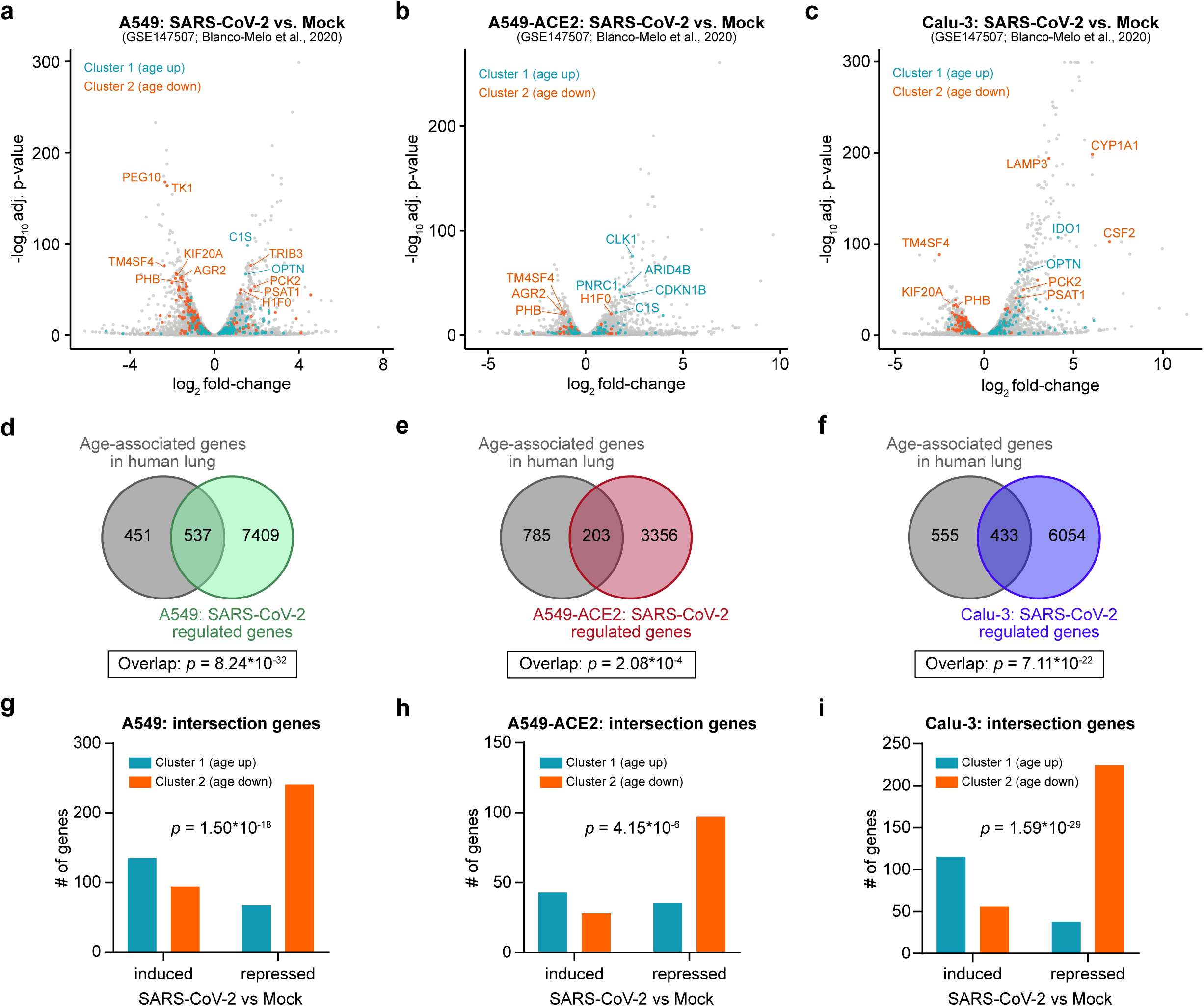
SARS-CoV-2 infection alters the expression of lung age-associated genes. **a-c**. Volcano plots of differentially expressed genes in A549 cells **(a)**, A549 cells transduced with an ACE2 vector (A549-ACE2) **(b)**, or Calu-3 cells **(c)**. Data are from GSE147507 (Blanco-Melo et al., 2020). Age-associated genes are color-coded. **d-f**. Venn diagrams highlighting the intersections between lung age-associated genes and SARS-CoV-2 regulated genes in A549 cells **(d)**, A549-ACE2 cells **(e)**, or Calu-3 cells **(f)**. Statistical significance of the overlap was assessed by hypergeometric test. **g-i**. Characteristics of age-associated genes that are affected by SARS-CoV-2 infection in A549 cells **(g)**, A549-ACE2 cells **(h)**, or Calu-3 cells **(i)**, from **d-f**. Statistical significance of the interaction between the directionality of SARS-CoV-2 regulation (induced or repressed) and the directionality of age-association (increase or decrease with age) was assessed by two-tailed Fischer’s exact test.

To identify a consensus set of age-associated genes that are regulated by SARS-CoV-2 infection, we integrated the analyses from all 3 cell lines. 603 genes were consistently induced by SARS-CoV-2 infection (**Figure 7a**). Of these, 20 genes are in Cluster 1 (increase with age) and 2 genes are in Cluster 2 (decrease with age). The 20 induced genes in Cluster 1 include several factors involved in RAS signaling (*RAB8B, RASA2*, and *RASGRP1*) as well as *CLK1*, which was shown to be involved in host responses to SARS-CoV infection (**Figure 4a-b**). On the other hand, 641 genes were concordantly repressed by SARS-CoV-2 infection (**Figure 7b**), with 7 genes in Cluster 1 and 49 genes in Cluster 2. Interestingly, ontology analysis of the 49 repressed genes in Cluster 2 revealed enrichment for mitochondrion, oxidoreductase, oxidative phosphorylation, pyruvate metabolism, and lysosome (**Figure 7c**), indicating that these pathways are affected both by aging and by SARS-CoV-2 infection.

**Figure 7:**
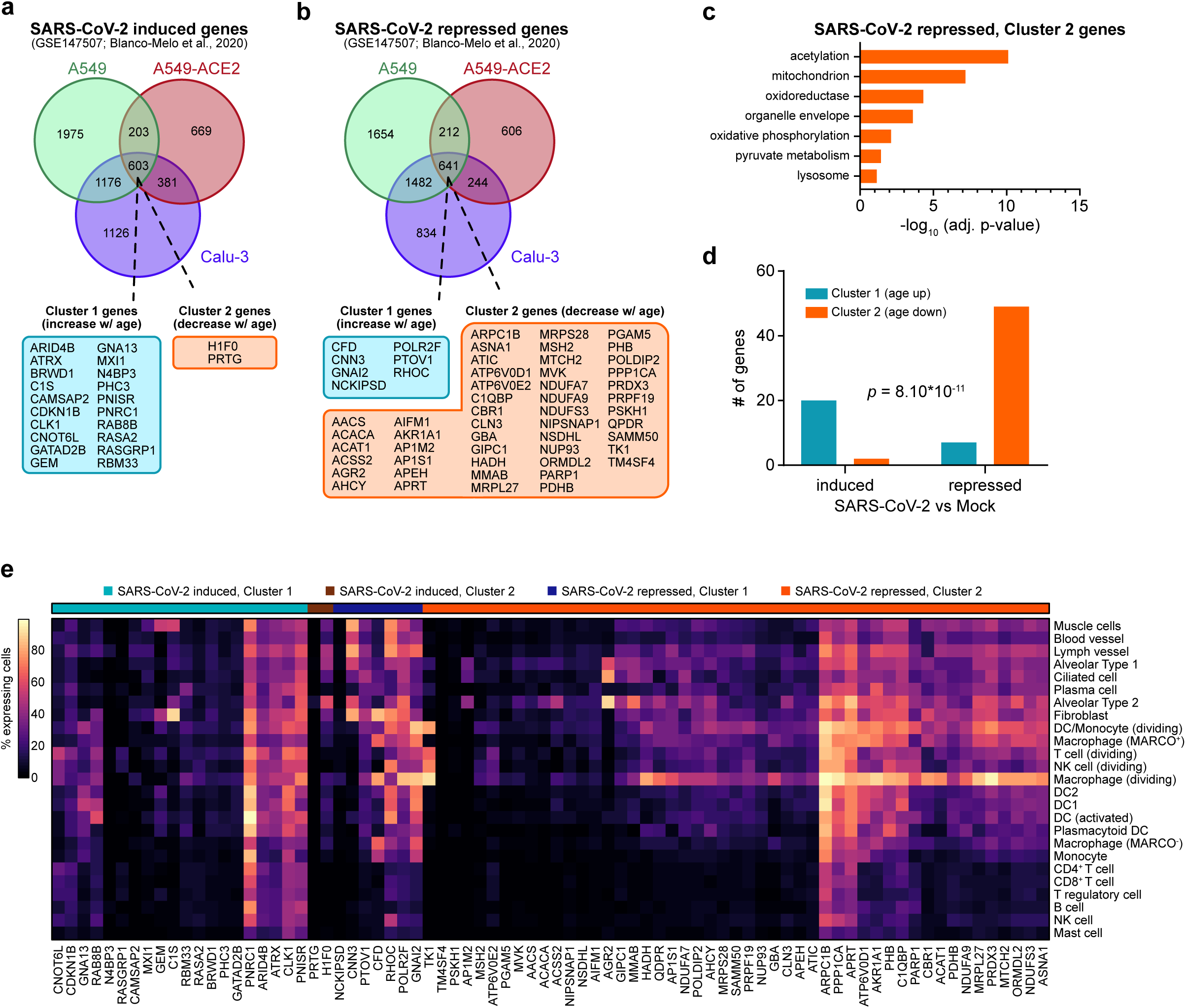
Transcriptional parallels between SARS-CoV-2 infection and the aging human lung. **a-b**. Venn diagram of shared SARS-CoV-2 induced genes **(a)** or SARS-CoV-2 repressed genes **(b)** in A549 cells, A549-ACE2 cells, and Calu-3 cells. Age-associated genes lying at the intersection of all 3 datasets are detailed, classified by their age-associated cluster. Data are from GSE147507 (Blanco-Melo et al., 2020). **c**. DAVID gene ontology and pathway analysis of Cluster 2 age-associated genes that are repressed upon SARS-CoV-2 infection. **d**. Characteristics of age-associated genes that are concordantly regulated by SARS-CoV-2 infection across all 3 cell lines, from **a-b**. Statistical significance of the interaction between the directionality of SARS-CoV-2 regulation (induced or repressed) and the directionality of age-association (increase or decrease with age) was assessed by two-tailed Fischer’s exact test. **e**. Heatmap showing the percentage of cells expressing each of the genes highlighted in **a-b**. Genes are annotated by whether they are induced or repressed by SARS-CoV-2 infection, and whether they increase or decrease in expression with age. Data are from the Tissue Stability Cell Atlas.

Within the consensus set of all 78 age-associated genes that are perturbed by SARS-CoV-2 infection, the directionality of SARS-CoV-2 regulation (induced or repressed) and the directionality of age-association (increase or decrease with age) were significantly associated (**Figure 7d**). Analysis of the human lung scRNA-seq datasets revealed the cell types that normally express these different genes (**Figure 7e** and **Supplementary Figure 7**). While many of these genes are broadly expressed across multiple cell types, some are preferentially expressed in AT2 cells (such as *PRTG, ACACA, ACSS2, AGR2, AP1M2*, and *MMAB*). Collectively, these analyses highlight the surprising parallels between the aging transcriptome of the human lung and the transcriptional changes caused by SARS-CoV-2 infection.

## Discussion

Here we systematically analyzed the transcriptome of the aging human lung and its relationship to SARS-CoV-2. We found that the aging lung is characterized by a wide array of changes that could contribute to the worse outcomes of older patients with COVID-19. On the transcriptional level, we first identified 1,285 genes that exhibit age-associated expression patterns. We subsequently demonstrated that the aging lung is characterized by several gene signatures, including increased vascular smooth muscle contraction, reduced mitochondrial activity, and decreased lipid metabolism. By integrating these data with single cell transcriptomes of human lung tissue, we further pinpointed the specific cell types that normally express the age-associated genes. We showed that lung epithelial cells, macrophages, and Th1 cells decrease in abundance with age, whereas fibroblasts and pericytes increase in abundance with age. These systematic changes in tissue composition and cell interactions can potentially propagate positive feedback loops that predispose the airways to pathological contraction ^69^. We find that some of the age-associated genes have been previously identified as host factors with a functional role in SARS-CoV replication ^63^, and a fraction of the age-associated factors have been shown to directly interact with the SARS-CoV-2 proteome ^64^. Furthermore, age-associated genes are significantly enriched among genes directly regulated by SARS-CoV-2 infection ^68^, suggesting transcriptional parallels between the aging lung and SARS-CoV-2 infection. Moreover, it is intriguing that the genes induced by SARS-CoV-2 infection tend to increase in expression with aging, and vice versa. Whether any of these age-associated changes causally contribute to the heightened susceptibility of COVID-19 in older populations remains to be experimentally tested.

It is also important to note that the datasets analyzed here were not from patients with COVID-19. Given the limited data that is currently publicly available, we emphasize that the analyses presented here at this stage should not be used to guide clinical practice. These analyses resulted in a number of previously unnoted observations and phenomena that illuminate new directions for subsequent research efforts on SARS-CoV-2, generating genetically-tractable hypotheses for why advanced age is one of the strongest risk factors for COVID-19 morbidity and mortality. Ultimately, we hope such knowledge can help the field to sooner develop rational therapies for COVID-19 that are rooted in concrete biological mechanisms.

## Acknowledgments

We thank Akiko Iwasaki, Craig Wilen, Hongyu Zhao, Wei Liu, Wenxuan Deng, Andre Levchenko, Katie Zhu, Ruth Montgomery, Bram Gerriten, Steven Kleinstein and a number of other colleagues for their critical comments and suggestions, which were incorporated into the analyses and manuscript. We thank Antonio Giraldez, Andre Levchenko, Chris Incarvito, Mike Crair, and Scott Strobel for their support on COVID-19 research. We thank our colleagues in the Chen lab, the Genetics Department, the Systems Biology Institute and various Yale entities. We also want to thank all of the healthcare workers who are risking their health on the frontlines to treat patients with this disease.

## Author contributions

RC and SC conceived and designed the study. RC developed the analysis approach, performed all data analyses, and created the figures. RC and SC prepared the manuscript. SC supervised the work.

---

**Supplementary Figure 1:**
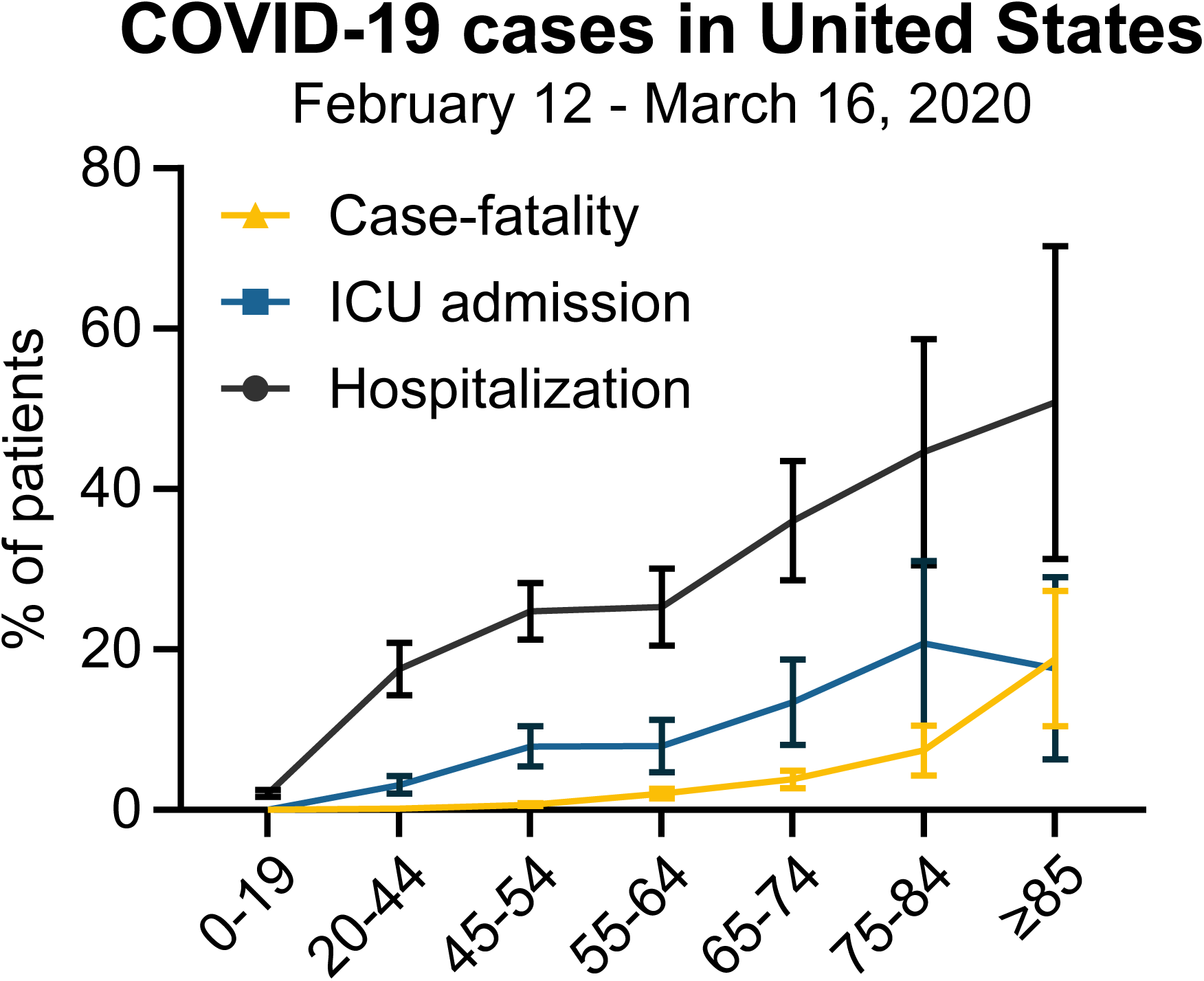
Severe outcomes of COVID-19 in the United States. Rates of hospitalization, ICU admission, and case-fatality from COVID-19 in the United States, stratified by age group. Data from the US CDC (as of March 16, 2020). Error bars represent lower and upper bounds of the estimated rates, given the preliminary nature of the data.

**Supplementary Figure 2:**
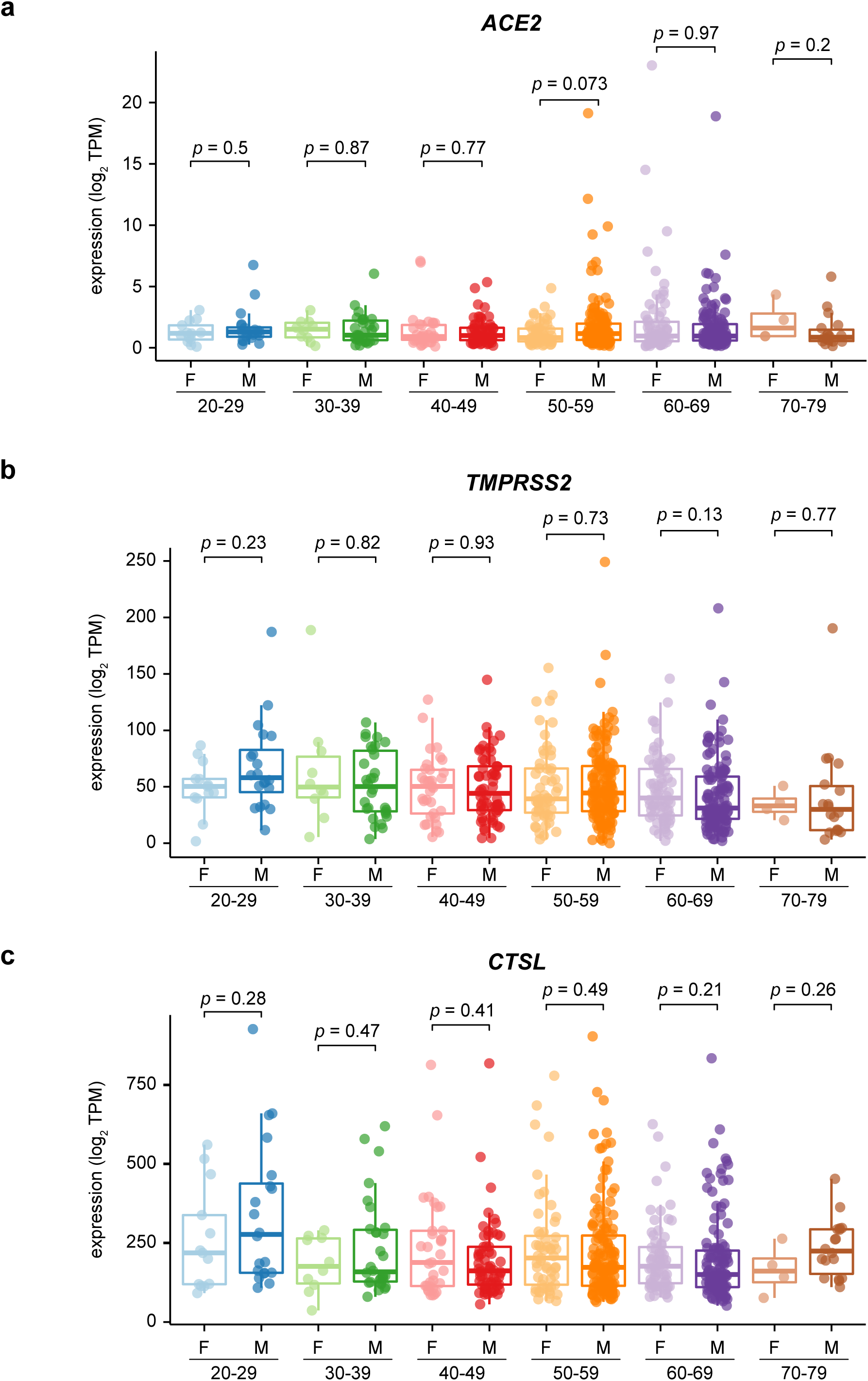
Sex-specific analysis of SARS-CoV-2 host entry factor expression. **a**. Expression of *ACE2* in lungs from female or male donors, stratified by age. Statistical significance was assessed by two-tailed Mann-Whitney test. **b**. Expression of *TMPRSS2* in lungs from female or male donors, stratified by age. Statistical significance was assessed by two-tailed Mann-Whitney test. **c**. Expression of *CTSL* in lungs from female or male donors, stratified by age. Statistical significance was assessed by two-tailed Mann-Whitney test.

**Supplementary Figure 3:**
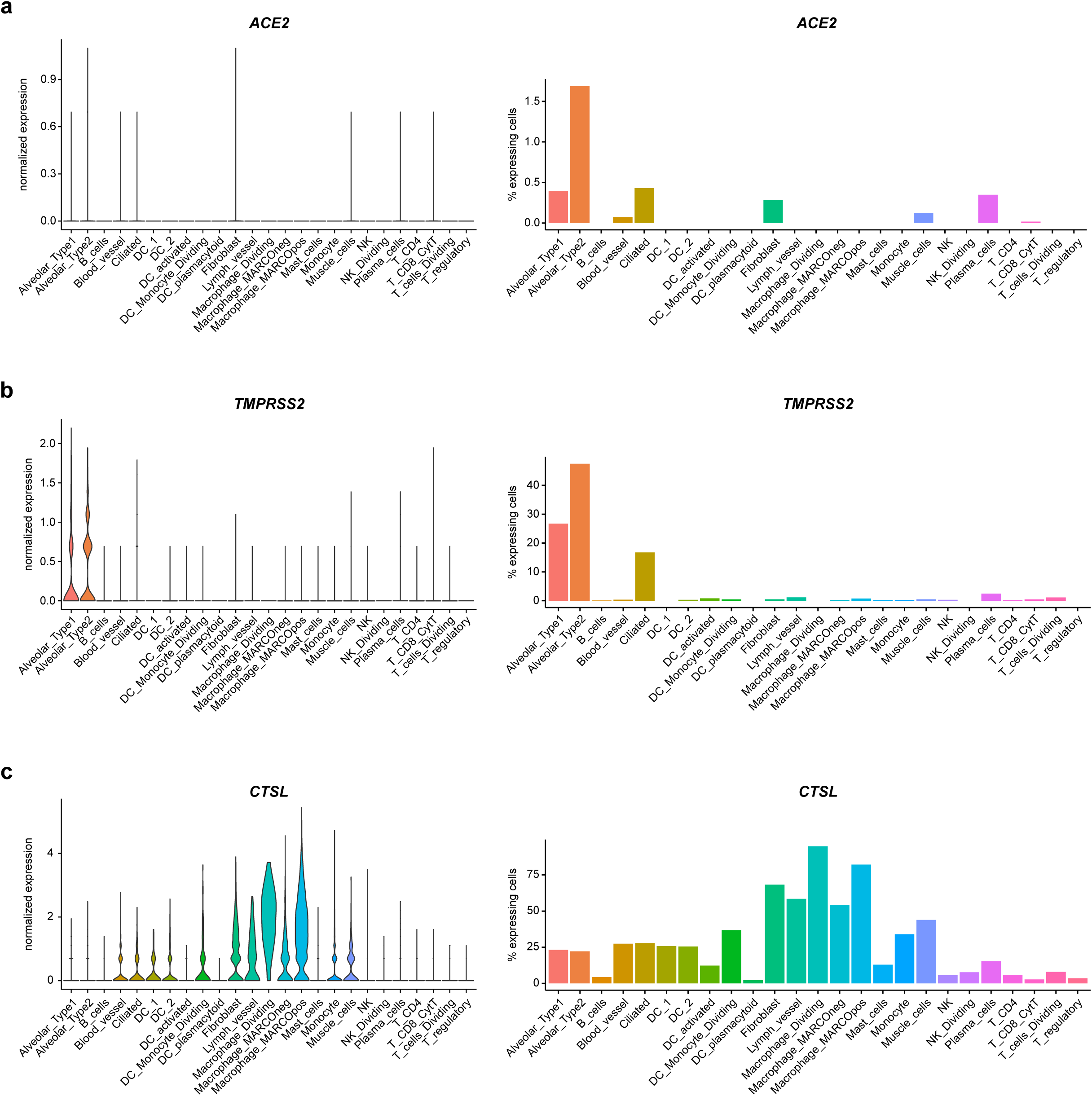
Expression of SARS-CoV-2 host factors across lung cell types. **a**. Left, normalized expression of *ACE2* across different lung cell types. Right, bar plot showing the percentage of *ACE2*-expressing cells, grouped by cell type. **b**. Left, normalized expression of *TMPRSS2* across different lung cell types. Right, bar plot showing the percentage of *TMPRSS2*-expressing cells, grouped by cell type. **c**. Left, normalized expression of *CTSL* across different lung cell types. Right, bar plot showing the percentage of *CTSL*-expressing cells, grouped by cell type.

**Supplementary Figure 4:**
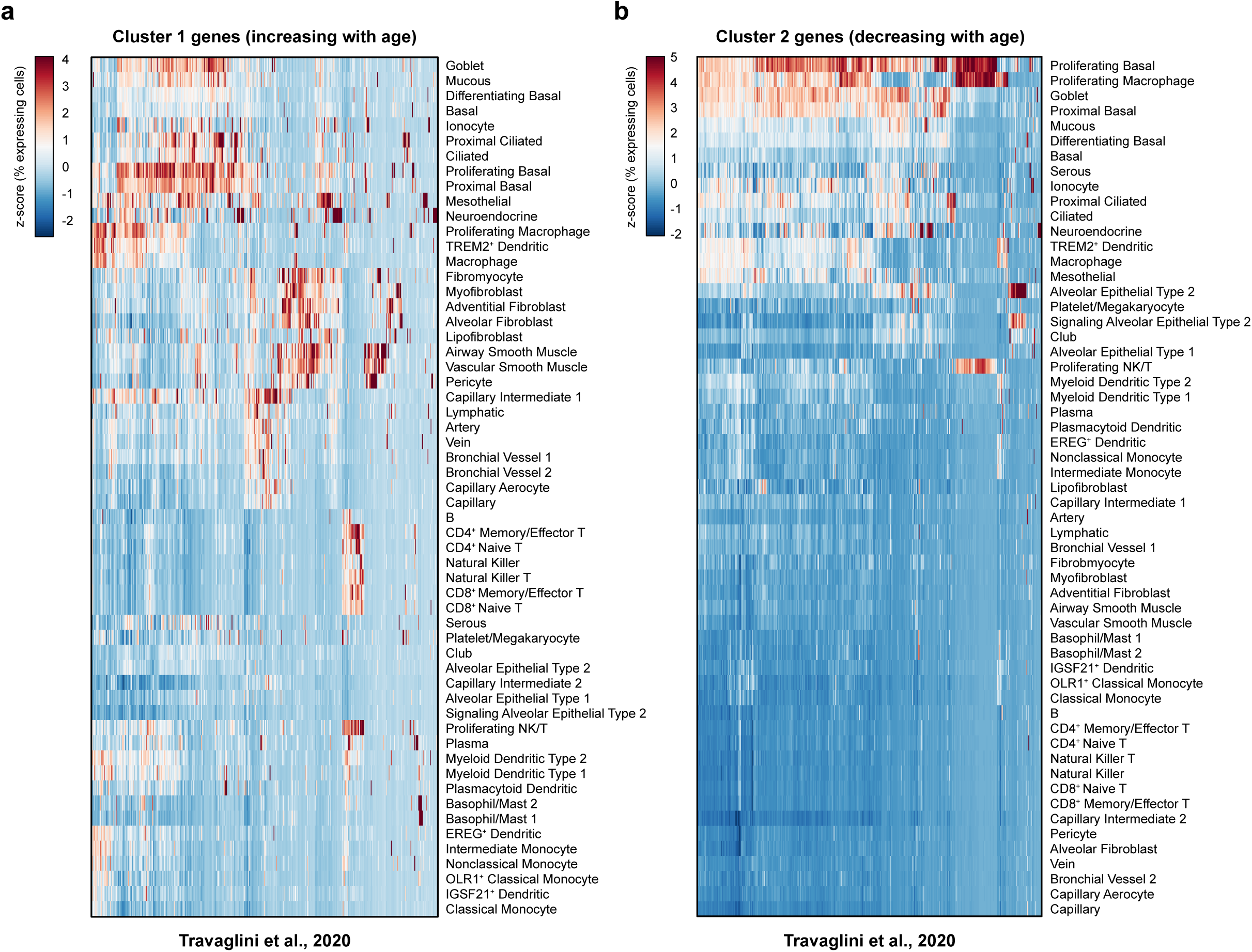
Additional cell type-specific analysis of age-associated genes. **a**. Heatmap showing the percentage of cells expressing each of the Cluster 1 genes (increasing with age), scaled by gene across the different cell types. Data are from the Human Lung Cell Atlas. **b**. Heatmap showing the percentage of cells expressing each of the Cluster 2 genes (decreasing with age), scaled by gene across the different cell types. Data are from the Human Lung Cell Atlas.

**Supplementary Figure 5:**
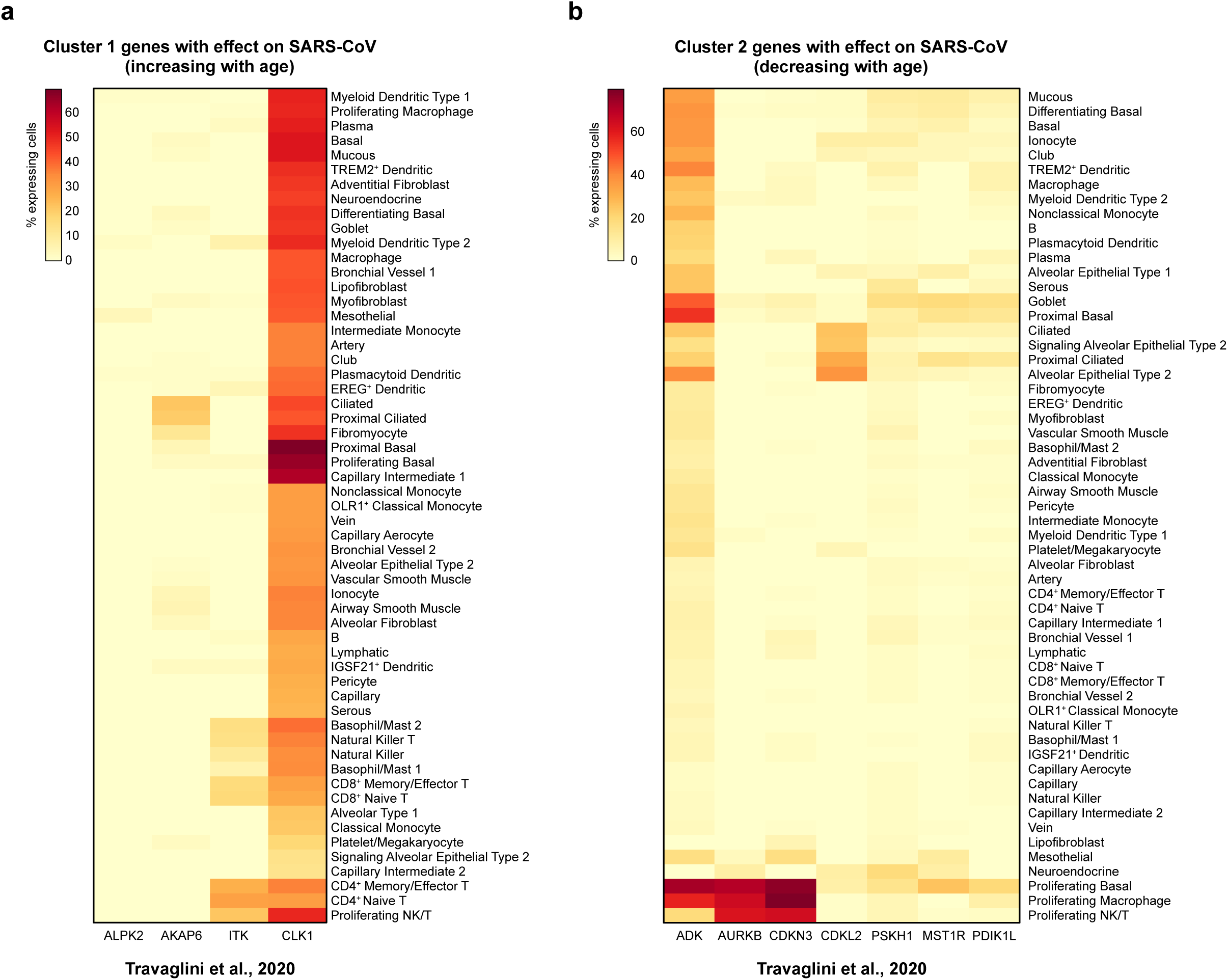
Additional cell type-specific analysis of age-associated genes that influence SARS-CoV replication. **a**. Heatmap showing the percentage of cells expressing each of the Cluster 1 genes (increasing with age) with an effect on SARS-CoV, as highlighted in **Figure 4a**. Data are from the Human Lung Cell Atlas. **b**. Heatmap showing the percentage of cells expressing each of the Cluster 2 genes (decreasing with age) with an effect on SARS-CoV, as highlighted in **Figure 4a**. Data are from the Human Lung Cell Atlas.

**Supplementary Figure 6:**
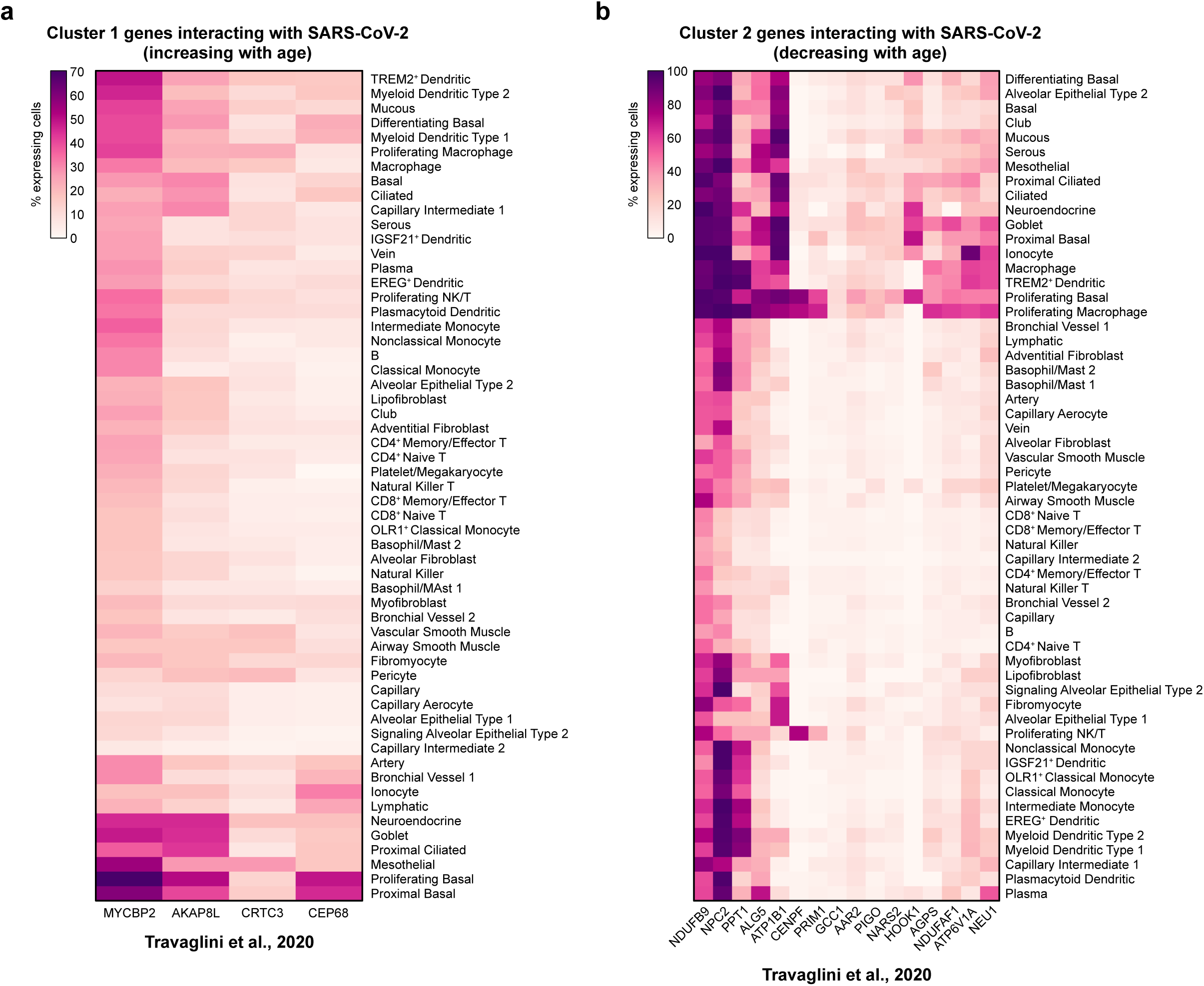
Additional cell type-specific analysis of age-associated factors that interact with SARS-CoV-2 proteins. **a**. Heatmap showing the percentage of cells expressing each of the Cluster 1 genes (increasing with age) that interact with SARS-CoV-2 proteins, as highlighted in **Figure 5b**. Data are from the Human Lung Cell Atlas. **b**. Heatmap showing the percentage of cells expressing each of the Cluster 2 genes (decreasing with age) that interact with SARS-CoV-2 proteins, as highlighted in **Figure 5b**. Data are from the Human Lung Cell Atlas.

**Supplementary Figure 7:**
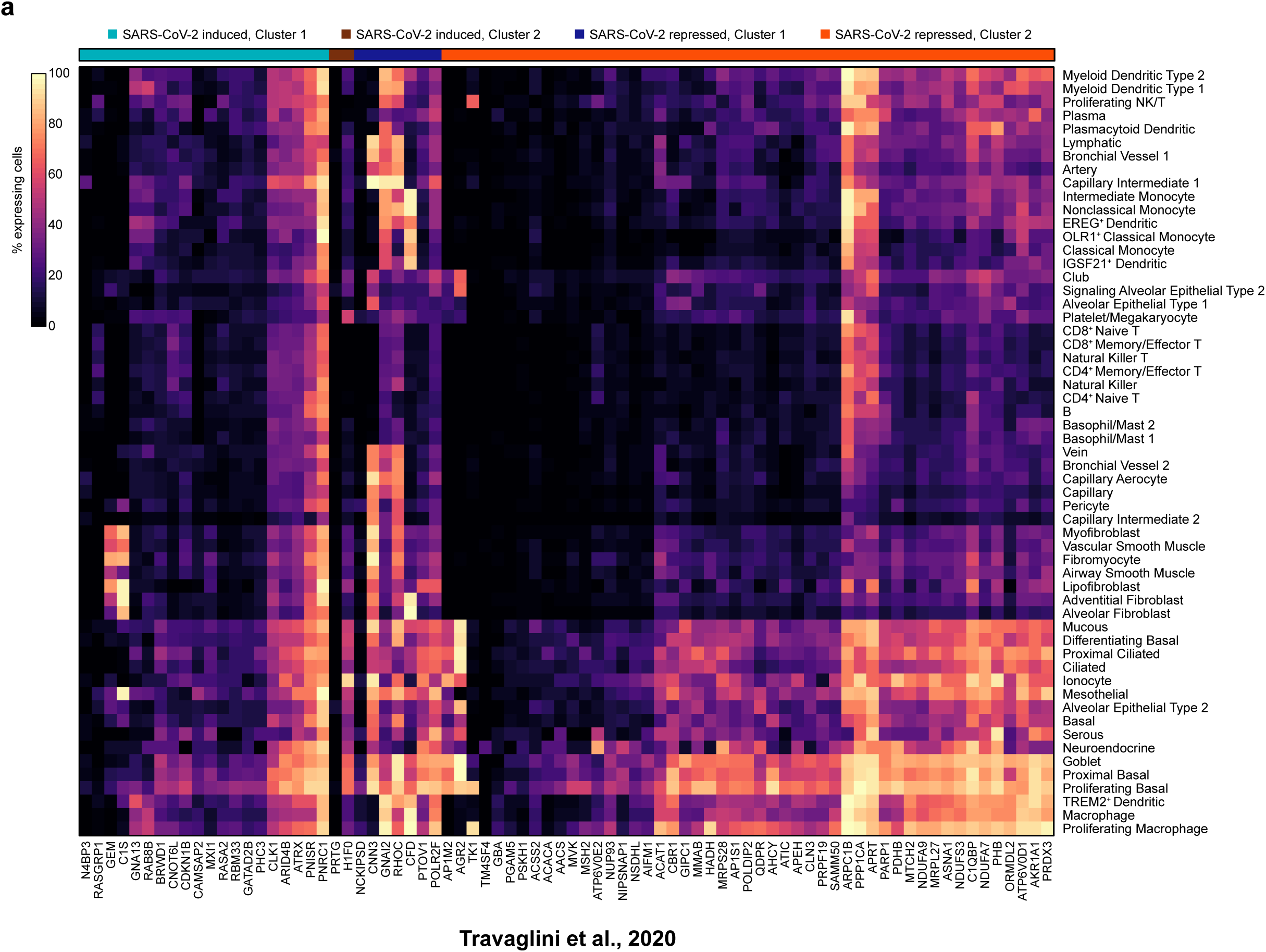
Additional cell type-specific analysis of age-associated factors that are differentially expressed upon SARS-CoV-2 infection. **a**. Heatmap showing the percentage of cells expressing each of the genes highlighted in **Figure 7a-b**. Genes are annotated by whether they are induced or repressed by SARS-CoV-2 infection, and whether they increase (Cluster 1) or decrease (Cluster 2) in expression with age. Data are from the Human Lung Cell Atlas.

## Methods

### Data accession

The Genotype-Tissue Expression (GTEx) Project was supported by the <u>Common Fund</u> of the Office of the Director of the National Institutes of Health, and by NCI, NHGRI, NHLBI, NIDA, NIMH, and NINDS ^10,11^. RNA-seq raw counts and normalized TPM matrices were downloaded from the GTEx Portal (https://gtexportal.org/home/index.html) on March 18, 2020, release v8. All accessed data used in this study are publicly available on the web portal and have been de-identified, except for patient age range and gender.

Case-fatality rates in China and Italy were from the Chinese CDC and Italian ISS, respectively, as described in a recent publication^3,4^. Statistics on COVID-19 cases in the United States from February 12 – March 16, 2020 were compiled from the CDC Morbidity and Mortality Weekly Report dated March 18, 2020 ^5^. Given the preliminary nature of the data, only estimated lower and upper bounds were reported.

Single cell transcriptomes of human lungs were obtained from the Tissue Stability Cell Atlas (https://www.tissuestabilitycellatlas.org/) ^19^ and from the Human Lung Cell Atlas (https://github.com/krasnowlab/HLCA and https://www.synapse.org/#!Synapse:syn21041850/) ^42^.

### RNA-seq gene expression visualization and statistical analysis

For visualization of RNA-seq expression data, the TPM values were log_2_ transformed and plotted in R (v3.6.1). All boxplots are Tukey boxplots, with interquartile range (IQR) boxes and 1.5 × IQR whiskers. Pairwise statistical comparisons in the plots were assessed by two-tailed Mann-Whitney test, while statistical comparisons across all age groups were performed by Kruskal-Wallis test.

### Identification of age-associated genes in human lung

To identify age-associated genes, the raw counts values were analyzed by DESeq2 (v1.24.0) ^22^, using the likelihood ratio test (LRT). Age-associated genes were determined at a significance threshold of adjusted *p* < 0.0001. Genes passing the significance threshold were then scaled to z-scores and clustered using the degPatterns function from the R package DEGreport (v1.20.0). Gene clusters with progressive and consistent trends with age were retained for downstream analysis.

### Gene ontology and pathway analysis of lung age-associated genes

Gene ontology and pathway enrichment analysis was performed using DAVID (v6.8) ^70^ (https://david.ncifcrf.gov/), separating the age-associated genes into the two clusters (increasing or decreasing with age), as described above.

### scRNA-seq data analysis

scRNA-seq data were analyzed in R (v3.6.1) using Seurat^71,72^ and custom scripts. Of the 1285 age-associated genes identified from GTEx bulk transcriptomes, 1049 genes were matched in the Tissue Stability Cell Atlas dataset and 1021 genes were matched in the Human Lung Cell Atlas dataset. To determine the percentage of cells expressing a given gene, the expression matrices were converted to binary matrices by setting a threshold of expression > 0. Cell type-specific expression frequencies for each gene were then calculated using the provided cell type annotations. To identify genes preferentially expressed in a specific cell type, we further scaled the expression frequencies in R to obtain z-scores. Data were visualized in R using the NMF package ^73^. Where applicable, gene ontology analysis was performed with DAVID (v6.8) ^70^, using genes with z-score > 2 in the cell type of interest for analysis.

### Inferring the cellular compositions of the human lung

To infer the cellular composition of each lung sample, we analyzed the TPM expression matrices using the xCell algorithm^45^. The resultant cell type enrichment tables were analyzed in R. For data visualization, cell type enrichment scores were scaled to z-scores, and the median z-score for each age group was expressed as a heatmap, using the superheat package ^74^. Age-association was assessed across all age groups by Kruskal-Wallis test.

### Assessing functional roles of lung age-associated genes in SARS-CoV

To assess whether any age-associated genes affect host responses to SARS-CoV (a coronavirus related to SARS-CoV-2), we analyzed the data from a published siRNA screen of host factors influencing SARS-CoV ^63^ (Data Set S1 in the publication; accessed on March 20, 2020). For data visualization, each point corresponding to a target gene was size-scaled and color-coded according to the age-association statistical analyses described above.

### Age-associated genes that interact with the SARS-CoV-2 proteome

To assess whether any lung age-associated genes encode proteins that interact with the SARS-CoV-2 proteome, we compiled the data from a preprint manuscript detailing the human host factors that interact with 27 different proteins in the SARS-CoV-2 proteome ^64^ (accessed on March 23, 2020).

### Age-associated genes that are transcriptionally regulated upon SARS-CoV-2 infection

To assess whether the expression of lung age-associated genes is influenced by SARS-CoV-2 infection, we utilized the data from a preprint manuscript detailing the transcriptional response to SARS-CoV-2 infection ^68^, from the Gene Expression Omnibus (https://www.ncbi.nlm.nih.gov/geo/query/acc.cgi?acc=GSE147507) (accessed on April 13, 2020). Differentially expressed genes were determined using the Wald test in DESeq2 (v1.24.0) ^22^ comparing SARS-CoV-2 infected cells to batch-matched mock controls, with a significance threshold of adjusted *p* < 0.05.

Of the 1285 age-associated genes, 988 genes were matched to the RNA-seq dataset. Statistical significance of overlaps between the gene sets was assessed by hypergeometric test, assuming 21,797 total genes as annotated in the RNA-seq dataset and 988 age-associated genes. Statistical significance of the association between the directionality of SARS-CoV-2 regulation and the directionality of age-association was assessed by two-tailed Fischer’s exact test. Gene ontology and pathway enrichment analysis was performed using DAVID (v6.8) ^70^ (https://david.ncifcrf.gov/).

### Statistical information summary

Comprehensive information on the statistical analyses used are included in various places, including the figures, figure legends and results, where the methods, significance, p-values and/or tails are described. All error bars have been defined in the figure legends or methods.

### Code availability

Codes used for data analysis or generation of the figures related to this study are available upon request to the corresponding author and will be deposited to GitHub upon publication for free public access.

### Data and resource availability

All relevant processed data generated during this study are included in this article and its supplementary information files. Raw data are from various sources as described above. All data and resources related to this study are freely available upon request to the corresponding author.

## Supplementary Tables

**Table S1:** Demographics of donors for GTEx lung samples.

**Table S2:** Normalized expression matrix for SARS-CoV-2 host entry factors in human lung.

**Table S3:** Cell type-specific expression frequencies for SARS-CoV-2 host entry factors in human lung (Tissue Stability Cell Atlas).

**Table S4:** Likelihood-ratio test table for evaluating age-associated genes.

**Table S5:** Final set of age-associated genes.

**Table S6:** DAVID analysis table for genes that increase in expression with age.

**Table S7:** DAVID analysis table for genes that decrease in expression with age.

**Table S8:** Cell type-specific expression frequencies for genes that increase in expression with age (Tissue Stability Cell Atlas).

**Table S9:** Cell type-specific expression frequencies for genes that increase in expression with age, transformed to z-scores (Tissue Stability Cell Atlas).

**Table S10:** Cell type-specific expression frequencies for genes that decrease in expression with age (Tissue Stability Cell Atlas).

**Table S11:** Cell type-specific expression frequencies for genes that decrease in expression with age, transformed to z-scores (Tissue Stability Cell Atlas).

**Table S12:** Cell type-specific expression frequencies for genes that increase in expression with age (Human Lung Cell Atlas).

**Table S13:** Cell type-specific expression frequencies for genes that increase in expression with age, transformed to z-scores (Human Lung Cell Atlas).

**Table S14:** Cell type-specific expression frequencies for genes that decrease in expression with age (Human Lung Cell Atlas).

**Table S15:** Cell type-specific expression frequencies for genes that decrease in expression with age, transformed to z-scores (Human Lung Cell Atlas).

**Table S16:** DAVID analysis table for muscle-enriched genes that increase in expression with age.

**Table S17:** DAVID analysis table for AT2-enriched genes that decrease in expression with age.

**Table S18:** Table of scaled cell enrichment scores, median value of each age group.

**Table S19:** Age-association statistics for cell enrichment scores.

**Table S20:** Annotation of a prior SARS-CoV siRNA screen (de Wilde et al., 2015) with age-association statistics from this study.

**Table S21:** Annotation of the SARS-CoV-2 : Human protein interactome (Gordon et al., 2020) with age-association statistics from this study.

**Table S22:** Differential expression analysis in A549 cells, infected with SARS-CoV-2 vs mock control, with age-association annotations.

**Table S23:** Differential expression analysis in A549-ACE2 cells, infected with SARS-CoV-2 vs mock control, with age-association annotations.

**Table S24:** Differential expression analysis in Calu-3 cells, infected with SARS-CoV-2 vs mock control, with age-association annotations.

**Table S25:** Consensus differentially expressed genes upon SARS-CoV-2 infection, intersected with age-associated genes.

**Table S26:** Characteristics of age-associated genes that are regulated by SARS-CoV-2 infection.

**Table S27:** DAVID analysis table for SARS-CoV-2 repressed genes that decrease in expression with age.

